# Sex-specific recombination predicts parent of origin for recurrent genomic disorders

**DOI:** 10.1101/2020.06.01.128553

**Authors:** Trenell Mosley, H. Richard Johnston, David J. Cutler, Michael E. Zwick, Jennifer G. Mulle

**Author notes:** Correspondence and request for materials should be addressed to Jennifer Gladys Mulle, MHS, PhD.

## Abstract

Genomic disorders are caused by structural rearrangements of the genome that generally occur during meiosis^1^. Often the rearrangements result in large-scale (> 1 kb) copy number variants (CNV; deletions or duplications ≥ 1 kb)^2,3^. Recurrent pathogenic CNVs harbor similar breakpoints in multiple unrelated individuals and are primarily formed via non-allelic homologous recombination (NAHR)^3,4^. Several pathogenic NAHR-mediated recurrent CNV loci demonstrate biases for parental origin of *de novo* CNVs^5–9^. However, the mechanism underlying these biases is not well understood. Here we have curated parent of origin data for multiple pathogenic CNV loci and demonstrate a significant association between sex-specific differences in meiotic recombination and parental origin biases at these loci. Our results suggest that parental-origin of CNVs is largely controlled by sex-specific recombination rates and bring into light the need to consider these differences when seeking to determine the factors underlying risk for structural variation.

## INTRODUCTION

Genomic disorders are caused by pathological structural variation in the human genome usually arising *de novo* during parental meiosis.^3,10–12^. The most common pathogenic variety of these rearrangements are copy number variants (CNVs), i.e. a deletion or duplication of > 1 kb of genetic material^11,13^. The clinical phenotypes of genomic disorders are varied. They include congenital dysmorphisms, neurodevelopmental, neurodegenerative and neuropsychiatric manifestations, and even more common complex phenotypes such as obesity and hypertension^14–19^. CNVs have been observed in 10% of sporadic cases of autism^20,21^, 15% of schizophrenia cases^22,23^, and 16% of cases of intellectual disability^24^. These and other associations highlight the importance of structural variation to human health and the need to understand the factors influencing how they arise.

There is an intense interest in understanding the mechanisms by which CNVs form^25,26^. In several regions of the genome, *de novo* CNVs with approximately the same breakpoints recur in independent meioses (recurrent CNVs)^3,4^. The presence of segmental duplications flanking these intervals is a hallmark feature of recurrent CNVs. It is hypothesized that misalignment and subsequent recombination between non-allelic low copy repeat (LCR) segments within the segmental duplication regions is the formative event giving rise to the CNV^27,28^, so called non-allelic homologous recombination (NAHR). Risk factors that may favor NAHR have been investigated and include sequence composition and orientation of the LCRs themselves^27,29^ as well as the presence of inversions at the locus^7,30^.

Parental bias for the origin of recurrent *de novo* CNVs remains unexplained. *De novo* deletions at the 16p11.2 and 17q11.2 loci are more likely to arise on maternally inherited chromosomes^5,8,31,32^. Deletions at the 22q11.2 locus show a slight maternal bias as well^6^. In contrast, deletions at the 5q35.3 locus (Sotos syndrome) display a paternal origin bias^9,33^. Deletions at the 7q11.23 locus (Williams syndrome) do not show a bias in parental origin^7^. While it has been suggested that sex-specific recombination hotspots might influence sex biases in NAHR^5^, this hypothesis has not been formally tested.

The majority of recurrent CNVs are thought to form during meiosis, when homologous chromosomes align and synapse during prophase I^34^. It is well established that meiosis differs significantly between males and females. In males, spermatagonia continuously divide and complete meiosis throughout postpubescent life with all four products of meiosis resulting in gametes. In contrast, in human females oogonia are established in fetal life and enter into a extended period of prolonged stasis in prophase I of meiosis until they complete meiosis upon ovulation and fertilization^35^. Additionally, in female meiosis, only one of four products of meiosis result in a gamete. Sexual dymorphism in meiosis extends to the patterns and processes of recombination during meiosis^34^. Here we seek to ask whether genome-wide sex-specific rates in meiotic recombination are coincident with bias in parent of origin for *de novo* CNV.

## RESULTS

### Recurrent Genomic Disorder Loci Literature Search

We conducted a systematic literature search for the 55 structural (Supplementary Data 1) variant loci in Coe et al., 2014^15^. We identified parent-of-origin studies that met the following criteria: (1) the study detailed parent of origin data for NAHR-mediated loci as designated by Coe et al., 2014^15^, (2) the study reported parent of origin data for non-imprinted loci, (3) the study reported data for more than 10 families with affected children, and (4) the study clearly treated monozygotic twins as one meiosis event. 25 studies met inclusion criteria; from these 25 studies, data were curated for six loci, including copy number variants at 5q35.3^9,36^, 7q11.23^1,7,37–42^, 16p11.2^5^, 17p11.2^43–45^, 17q11.2^8,31,32^, and 22q11.2^6,38,46–51^ (Table 1). Each locus has between one and eight independent studies representing in total 1,438 *de novo* deletion and duplication events.

**Table 1.** Summary of genomic disorder loci and calculated linear regression variables

| Locus | CNV Type | OMIM Number | BED Coordinates<sup>15</sup> | Total Samples | *M:F Origin Ratio | <sup>†</sup>Log<sub>e</sub> M:F Recombination Ratio<sup>52</sup> | <sup>‡</sup>Study Refs. |
| --- | --- | --- | --- | --- | --- | --- | --- |
| 3q29 | Del | 609425 | chr3:195988732-197628732 | 14 | 13:1 | 2.4239321 | Current Study |
| 5q35.3 | Del | 117550 | chr5:176290391-177630393 | 41 | 36:5 | 0.2828221 | 9,33 |
| 7q11.23 | Del/Dup | 194050/ 609757 | chr7:73328061-74727726 | 530 | 251:279 | -1.3620333 | 1,7,37–42 |
| 16p11.2 | Del/Dup | 611913/ 614671 | chr16:29641178-30191178 | 79 | 8:71 | -2.9835893 | 5 |
| 17p11.2 | Del | 182290 | chr17:16805961-20576095 | 59 | 34:25 | -1.8419594 | 43–45 |
| 17q11.2 | Del | 162200 | chr17:30838856-31888868 | 73 | 12:61 | -1.9637162 | 8,31,32 |
| 22q11.2 | Del | 188400/192430 | chr22:19032487-20302477 | 642 | 275:367 | -0.9630031 | 6,38,46–51 |

\*Male to female CNV parent of origin count ratio
<sup>†</sup>Natural log-transformed male to female recombination rate ratio
<sup>‡</sup>Independent studies from which the parent of origin data for the current analysis were obtained.
Summary of loci analyzed. BED coordinates correspond to Hg38 (LiftOver from hg19 coordinates in Coe et al., 2014<sup>15</sup>). Log<sub>e</sub>-transformed male to female recombination rate ratios are calculated from recombination rates (cM/Mb) reported in Halldorsson et al., 2019<sup>52</sup>.

### Parent of Origin of 3q29 Deletion

We determined parent of origin in 12 full trios where a proband had a *de novo* 3q29 deletion; in 2 additional trios where only proband and maternal DNA samples were available, parent of origin was inferred. In the 14 trios, 13 deletions (92.9%) arose on the paternal genome indicating a significant departure from the null expectation of 50% (*p* = 0.002, binomial exact). When accounting for only full trios, 11 of 12 (91.7%) deletions arose on paternal haplotypes (*p* = 0.006, binomial exact), altogether indicating there is a paternal bias for origin of the 3q29 deletion (Supplementary Table 1). We examined the age distribution of male parents of origin in our cohort. The mean age of the male parents of origin in our cohort is 34 years (median = 34 years), and is not significantly different from the 2018 U.S national average, (31.8 years) (*p* = 0.08, Two-tailed two sample t-test,) indicating the bias is unlikely to be due oversampling of older fathers (Supplementary Table 1).

### Meiotic Recombination and Parental Origin

For each locus, we addressed the possibility that meiotic recombination could explain the parental biases observed for recurrent genomic disorder loci mediated by NAHR. We calculated the average male and female recombination rates (cM/Mb) across each critical region (as cited in Coe et al., 2014^15^) for all seven loci using recombination rates published by the deCODE genetics group^52^. We regressed the combined male to female parental origin count ratios for each locus on the male to female recombination rate ratio (Table 1; Supplementary Table 2). Recombination rate ratios explain 83.44% of the variation in male to female parental origin of CNVs (multiple R^2^ = 0.8344; *p* = 0.004) (Figure 1). This estimate is not influenced by any particular data point as demonstrated by a sensitivity analysis (Supplementary Table 3). Additionally, separating deletions and duplications to account for potential differences in the distributions of parental origins produced similar results (multiple R^2^ = 0.8338; *p* = 0.0005) (Supplementary Figure 1; Supplementary Table 4). This analysis reveals that sex-specific differences in recombination rates predict parental origin of genomic disorder CNVs.

**Figure 1.**
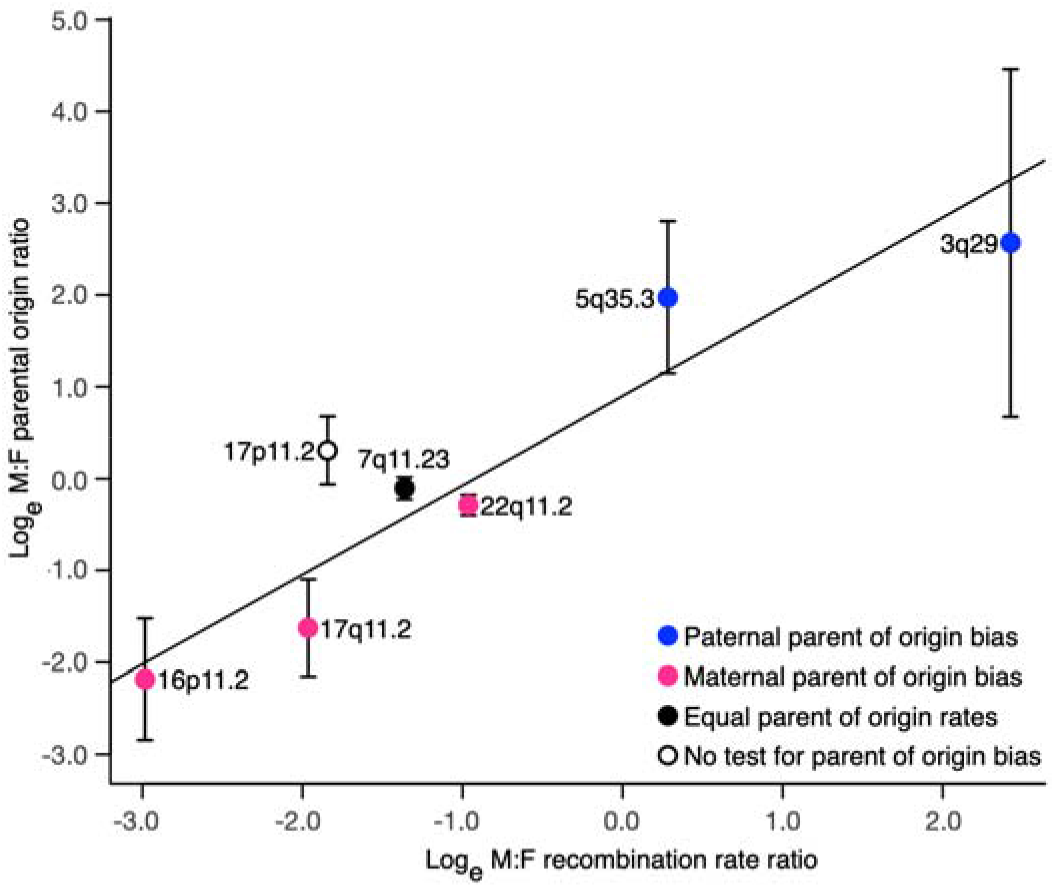
Recombination rates associate with parental origin bias. Log_e_-transformed male to female parental origin ratio regressed on log_e_-transformed male to female recombination rate ratio. Recombination rates are associated with male-to-female parental origin ratios (mulitple R^2^ = 0.8344, *p*=0.004, β = 0.9728, 95% CI: (0.47449531,1.471012)). Curated parent of origin data from multiple published studies is collapsed by loci into single data points. Loci are color coded by reported bias. Blue: loci with reported paternal biases, pink: loci with maternal biases, black: loci with equal male and female parental origin rates, white: loci where no test for parental origin bias conducted in literature. Error bars correspond to upper and lower bounds of 95% confidence interval.

## DISCUSSION

Parent of origin bias for *de novo* events at recurrent CNV loci has lacked a compelling explanation. Our analysis of data gathered from 25 published parent of origin reports demonstrate that sex-specific variation in local meiotic recombination rates explains approximately 83% of parent of origin bias at recurrent CNV loci. Human male and female meiotic recombination rates and patterns differ greatly across the broad scale of human chromosomes. Recombination events are nearly uniformly distributed across the chromosome arms in females but tend to be clustered closer to the telomeres in males^53^. Our analysis reveals a parallel trend, such that CNVs contributing to genomic disorders arising more frequently in female meiosis occur closer to the centromeres of their respective chromosomes, while those exhibiting paternal biases occur closer to the telomeres. We note that this pattern has been previously recognized^5^. Here we have formally tested the hypothesis that recombination variation drives parent of orgin variantion in a rigorus, statistical framework and provided an estimate for the variance in parent of origin bias that is due to sex-dependent recombination rates.

Investigations into the mechanism by which recurrent CNVs arise have focused on LCRs and their makeup^3,54^. These regions are composed of units of sequence repeats that vary in orientation, percent homology, length, and copy number. Consequently, LCRs are mosaics of varying units, imparting complexity to LCR architecture^29^. The frequency of NAHR events mediated by LCRs is a function of these characteristics, and other features of the genomic architecture^30^. Specifically, the rate of NAHR is known to correlate positively with LCR length and percent homology and decrease as the distance between LCRs increases^26,27^. However, because LCRs are challenging to study with short-read sequencing technology, the population-level variability of these regions is not well described^55^. Recent breakthroughs with long-read sequencing and optical mapping have revealed remarkable variation in LCRs^56–58^, and haplotypes with higher risks for CNV formation have now been identified^59^. Our data suggest that any evaluation of CNV formation would be well served to consider the local meiotic recombination landscape. LCRs are substrates for NAHR^3^, and thus are subject to the recombination process. Local recombination rates may influence how likely an NAHR event will happen between two LCRs. Therefore, when analyzing LCR haplotypes and their susceptibility to NAHR, one would need to take into account sex-differences in recombination. For example, at loci with maternal biases, specific risk haplotypes may be required for males to form CNVs and vice versa. Greater enrichment of GC content, homologous core duplicons or the PRDM9 motifs, or other recombination-favoring factors may also be required^3,26^

While variation in recombination rates between sexes is well established^53,60–63^, prediction of individual risk may also need to consider individual variation in meiotic recombination, which is itself a heritable trait^61^. Here we show that 83% of the variation is explained by *mean* recombination rates in males and females. It could be that the remaining 17% is explained by individual level variation in rates. Variants in several genes, including *PRDM9,* have been shown to affect recombination rates and the distribution of double-stranded breaks in mammals^64,65^. Common alleles in *PRDM9* are evidenced to affect the percentage of recombination events within individuals that take place at hotspots^65^. Additionally, evidence shows that sex-specific hotspots exist in the genome and coincide with CNV loci^5,61^. While CNVs at 22q11.2 shows a slight maternal bias^6^, the maternal bias evident at 16p11.2 bias is relatively more apparent^5,6^. This may be due to the existence of a female hotspot at the 16p11.2 locus^5^. Existence of sex-specific hotspots may influence the likelihood of a recombination event in NAHR-prone regions in a particular sex and influence the strength of the parental bias in regions.

Many human genetic studies have observed correlations between inversion polymorphisms and genomic disorder loci^30,66^. Because these inversions are copy-number neutral and often located in complex repeat regions,^67^ they can be difficult to assay with current high-throughput strategies and their true impact remains to be explored. One model proposes that during meiosis these regions may fail to synapse properly and increase the probability of NAHR^68,69^. Another theory suggests formation of inversions increase directly oriented content in LCRs leading to a NAHR-favorable haplotype^70^. Supporting these theories, inversion polymorphisms have been identified at the majority of recurrent CNV loci^6,7,30,66,68,70,71^. At the 7q11.23, 17q21.31, and 5q35.3 loci^7,30,71^, compelling data indicts inversions as a highly associated marker of CNV formation. However, heterozygous inversions are known to *suppress* recombination perturbing the local pattern of recombination and altering the fate of chiasmata^72^. The analysis presented here strongly suggests that recombination is the driving force for CNV formation giving rise to an alternate explanation for the association between inversions and CNVs; They are both the consequence (and neither one the cause) of recombination between non-allelic homologous LCRs. Inversions and CNVs appear to be associated because both are being initiated by aberrant recombination. Viewing the system in this manner also explains the frequency of individual inversions at CNV loci. Inversions are arising via rare aberrant recombination, like CNVs, but subsequently being driven to higher frequency by natural selection, because they act to suppress recombination and “save offspring” from deleterious genomic disorders. Of course, frequent mutations leading to inversions and the details of LCR structure such as relative orientation and homology within a genomic region may promote or impede CNV formation in a locus-specific manner^73–75^. Further exploration of this relationship with improved genomic mapping can test these alternative models^76^. One testable prediction of the model described here is that inversions should be at higher frequency at loci giving rise to highly deleterious CNVs, as opposed to loci harboring recurrent benign CNVs.

To our knowledge, this study is the first comprehensive investigation of parental origin of NAHR-mediated CNV loci. Hehir-Kwa et al., and Ma et al., conducted similar large-scale studies, focusing on intellectual disability, developmental delay and congenital dysmorphisms, and determined a paternal bias for a sample predominantly of non-recurrent CNV^77,78^. They hypothesized that replication-based mechanisms of CNV formation contributed to the bias. Our study focuses on loci predicted to be formed via NAHR, and thus isolates our data from confounding by multiple mechanisms of CNV formation. Although our analysis includes data from over 1,400 samples, it is limited to existing studies on pathogenic rare CNVs. It does not include benign CNVs such as the 7q11.2 deletion^79^, since parent of origin data is scarce for these non-pathogenic loci. Analysis of a larger cohort of CNV loci including benign CNVs will give greater insight into the role of recombination, and sex differences in recombination in influencing parent of origin in CNVs.

Our data show that meiotic recombination predicts the parent of origin for recurrent CNVs underlying genomic disorders. The influence of recombination in CNV formation may also influence the incidence of recurrent CNVs. Females, on average, have more crossovers per genome, and when observing the frequency of CNVs in the population with known parental biases, a pattern emerges. 22q11.2 and 16p11.2 have a maternal bias and a prevalence of 1 in 4,000 and 1 in 3,000^6,80^ respectively, while 3q29 and 5q35.3 have paternal biases and a prevalence of 1 in 30,000 and 1 in 14,000 respectively^81^. While prevalence could be confounded by severity of the disorders, our data suggest the sex specific frequency of meiotic recombination may also influence the incidence of these genomic disorders. Investigation into this possibility could be achieved with a cohort of individuals with benign CNVs to reduce confounding by severity. Combining the sex-specific recombination landscape and the mechanistic factors underlying it with a more detailed understanding of existing structural factors at genomic disorder loci can be expected to help guide standards used to identify and perform genetic counseling for individuals at risk of genomic rearrangement.

## SUBJECTS AND METHODS

### Parent of Origin Determination

#### Literature Search and Data Curation

A set of known genomic disorder CNV loci mediated by non-allelic homologous recombination (NAHR) was curated from Coe et al., 2014^15^. This paper is an expansion of Cooper et al., 2011^82^ and includes breakpoint coordinates, syndrome name (if applicable), and whether the loci are flanked by LCRs, *i.e.* mediated by NAHR. Studies that reported parental origin of CNV (deletions and duplications) at these loci were curated using a systematic PubMed search and a set of inclusion criteria (Supplementary Methods and Supplementary Data 1). Seven loci were subsequently used for further analysis on the basis of the criteria mentioned above: 3q29 (current study), 5q35.3^9,33^, 7q11.23^1,7,37–42^, 16p11.2^5^, 17p11.2^43–45^, 17q11.2^8,31,32^, 22q11.2^6,38,46–51^.

#### Determination of Parental Origin for 3q29 Deletion

##### Study Subject Recruitment

Individuals with a clinically confirmed diagnosis of 3q29 deletion were ascertained through the internet-based 3q29 registry (https://3q29deletion.patientcrossroads.org/) as previously described^83^. We obtained blood samples and determined parental origin of the 3q329 deletion in 14 families. SNP genotyping was performed on 12 of the 14 families (10 full trios, 2 mother-child pairs) using the Illumina GSA-24 v 3.0 array. For 2 full trios (6 samples), parent of origin was determined from whole genome sequence data on Illumina’s NovaSeq 6000 platform. Quality control was performed with PLINK 1.9^84^ and our custom pipeline (Supplementary Table 1; See Supplementary Methods for more details).

##### Parental Origin Analysis

Parental origin of the 3q29 deletion was determined using PLINK 1.9 SNPs located within the 3q29 deletion critical region (chr3:196029182-197617792; hg38) were isolated for analysis. Mendelian errors (MEs) were called. The parent with the most MEs was considered the parent of origin for the 3q29 deletion (See Supplementary Methods for more details).

### Calculation of Recombination Rates

Chromosome male and female recombination rates (cM/Mb) were obtained from the deCODE sex-specific maps^52^. The data from deCODE is presented as binned rates across separate chromosomes. As such, each binned recombination rate was weighted by the total base pairs of CNV contained within the respective bin (breakpoints cited in Coe et al, 2014^15^). Weighted binned rates were then averaged across the CNV interval.

### Linear Regression Analysis

Parental origin data from multiple studies for each CNV locus were combined into one data point per locus after it was determined samples did not overlap between studies. The combined log_e_-transformed male-to-female parental origin count ratios for each locus was regressed on the calculated log_e_-transformed average male to female recombination rate ratio for that locus’ CNV interval. Each locus was weighted based on its combined sample size. Under the assumption that an NAHR event produces reciprocal deletion and duplication products, formation of both types of CNVs would be subject to the same mechanistic forces. Thus, for each locus, duplications and deletions were treated equally and grouped under one locus. See Table 1 and Supplementary Table 2 for the data calculated and used in the weighted linear regression.

## Data Availability

The data that support the findings of this study are available from the corresponding author upon reasonable request.

## Code Availability

The custom code that supports the findings of this study are available from the corresponding author upon reasonable request.

## ACKNOWLEDGEMENTS

The authors gratefully acknowledge the contributions of the members of the Emory 3q29 Project: Jennifer Mulle, Hallie Averbach, Katrina Aberizk, Emily Black, Gary J. Bassell, T. Lindsey Burrell, Grace Carlock, Shanthi Cambala, Tamara Caspary, Joseph F. Cubells, David Cutler, Paul A. Dawson, Michael P. Epstein, Roberto Espana, Michael J. Gambello, Katrina Goines, Sandra M. Goulding, Ryan Guest, Henry R. Johnston, Cheryl Klaiman, Sookyong Koh, Elizabeth J. Leslie, Longchuan Li, Bryan Mak, Tamika Malone, Michael Mortillo, Trenell Mosley, Melissa M. Murphy, Derek Novacek, Becky Pollak, Ryan Purcell, Timothy Rutkowski, Rossana Sanchez, Celine A. Saulnier, Jason Schroeder, Esra Sefik, Brittney Sholar, Sarah Shultz, Nikisha Sisodiya, Steven Sloan, Elaine F. Walker, Stephen T. Warren, David Weinshenker, and Zhexing Wen, Michael C. Zinsmeister, Michael E. Zwick. This study was supported in part by the Emory Integrated Genomics Core (EIGC), which is subsidized by the Emory University School of Medicine and is one of the Emory Integrated Core Facilities. Additional support was provided by the Georgia Clinical & Translational Science Alliance of the National Institutes of Health under Award Number UL1TR002378. The content is solely the responsibility of the authors and does not necessarily reflect the official views of the National Institutes of Health. Funding support includes Emory University, Grant/Award Number: Treasure Your Exceptions Project; National Institutes of Health, Grant/Award Numbers: 1R01MH110701□01A1, F31GM131609, T32 GM0008490.

## AUTHOR CONTRIBUTIONS

All authors assisted with data analysis, hypothesis generation, and writing the manuscript.

## ADDITIONAL INFORMATION

### Ethics Approval and Consent to Participate

This study was approved by Emory University’s Institutional Review Board (IRB00064133). All study subjects gave informed consent prior to participating in this study.

## Supplementary Information

Supplementary information is available for this paper.

