## Supplementary Materials for "Sex-specific recombination predicts parent of origin for recurrent genomic disorders"

### SUPPLEMENTARY INFORMATION

- Supplementary Methods
- Supplementary References
- Supplementary Figure 1
- Supplementary Table 1
- Supplementary Table 2
- Supplementary Table 3
- Supplementary Table 4

### SUBJECTS AND METHODS

#### Literature Search and Data Curation

Genomic disorder CNV loci mediated by NAHR and breakpoint coordinates were curated from Coe et al., 2014[1]. This paper is an expansion of Cooper et al., 2011[2]. Parental origin data for the CNV (deletions and duplications) at these loci were curated using a systematic PubMed search:

On PubMed loci were searched using a phrase with the format: *cytogenic locus OR syndrome.* The number of results/hits was recorded and if the initial search produced more than 100 hits, a sub-search was performed. This sub-search used the following format: *(cytogenic locus OR syndrome)* + *(parental bias OR parental origin OR transmission bias OR parent-of-origin OR parent of origin OR maternal bias OR paternal bias).*

Studies were excluded from the analysis:

- If the study reported parental origin for less than 10 families
- If the locus under investigation is subject to genomic imprinting
- If it was unclear if the study treated monozygotic twins as one or two transmissions in the analysis

All search phrases and studies curated as part of the current analysis and the loci used in the linear regression analysis are listed in Supplementary Table S1, and Table 1 & Extended Data Table 2, respectively.

**Study Subject Recruitment**

Individuals with clinically confirmed diagnosis of 3q29 deletion were ascertained through the internet-based 3q29 registry (https://3q29deletion.patientcrossroads.org/) as previously described[3,4]. We obtained blood samples and determined parental origin of the 3q29 deletion in 14 families. Of the 14 families, 12 were full trios. The remaining two families were both mother and child pairs.

**Sample Collection and Banking**

Whole blood was collected from proband, mother and father as previously reported[4] and banked at the NIMH Repository and Genomics Resource (NRGR; Piscataway, New Jersey, USA).

**DNA Isolation**

DNA samples were isolated from/obtained from biobanked samples at the NIMH Repository and Genomics Resource (NRGR; Piscataway, New Jersey, USA). The source of DNA was either whole blood or LCLs derived from biobanked blood samples.

**SNP Genotyping and QC**

SNP genotyping was performed on 12 of the 14 families (10 full trios, 2 mother-child pairs) by AKESOgen (Peachtree Corners, Georgia, USA) on the Illumina GSA-24 v 3.0 array. DNA from participants was normalized genotyped according to AKESOgen/Illumina protocols. Data was returned as separate final reports that were combined into one final report, deduplicated and converted into PLINK format for quality control. QC was performed with PLINK 1.9[5]. Briefly, unmappable SNPs and SNPs with call rates less that 97% were dropped from the SNP call set. Samples reported sex and family relationships were verified using this set of quality SNPs and PLINK 1.9[5]. F coefficient estimates for the X chromosome were calculated and sex assignment was inferred for each sample in the batch. Before sex was inferred, the *--split-x* flag with the *hg38* modifier was used to identify pseudoautosomal regions of the X chromosomes for subsequent removal during the sex check. The default parameters for the *--check-sex* flag were used to infer sex. A sample with an F coefficient less than or equal to 0.2 was assigned as male, and a sample with an F coefficient greater than or equal to 0.9 was assigned as female. Any samples with an opposite sex assignment than indicated by the given pedigree were flagged and investigated for possible sample swapping or sample mixture. Expected relationships between related samples of the batch were verified with PLINK 1.9[5]. Variants were LD-pruned using the *--indep-pairwise* flag using 50, 5, and 0.2 for the variant count window size, variant count step size and $r^2$, respectively. The *--genome* flag was used to infer relationships (coefficient of relatedness; *r*) on this set of pruned SNPs. Among the control genomes there was a known parent-offspring relationship, which was used as a positive control, while the remaining control genomes were known to be unrelated. Any samples with an *r* estimate that indicated a degree of relationship (1st, 2nd, 3rd, etc.) different from the expected degree of relationship were flagged and investigated for possible sample swapping, sample mixture, or misinformation. All samples sex information and relationship information were concordant with our expectation based on information provided by the families.

**Whole-genome Sequencing**

For 2 full trios (families 3206 and 3147; 6 samples), parent of origin was determined from whole genome sequence data. All samples were sequenced at the Hudson–Alpha Institute of Biotechnology (Birmingham, Alabama, USA) using their published protocols. Sequencing was performed to approximately 30X coverage per genome on the Illumina NovaSeq 6000 platform. Following sequencing all base-calling was performed using standard Illumina software to generate the final FASTQ files for each sample.

**Sequence Alignment: PEMapper[6]**

FASTQ files were aligned on a per sample basis with PEMapper[6] using default parameters and a Smith-Waterman alignment threshold of 95%, as recommended for 150-bp paired-end reads. Alignment was performed relative to the human Hg38 reference as reported by the University of California at Santa Cruz (UCSC) Genome Browser on July 1, 2015. The output from PEMapper[6], pileup and indel files, were used as input for variant calling with PECaller[6]. Pileup files contained the number of reads where an A, C, G, or T nucleotide was seen together with the number of times that base appeared deleted or there was an insertion immediately after the base. Indel files contained the nucleotide sequence of the deletions and insertions indicated in the pileup files. Alignment performance was checked before moving to variant calling. Any sample(s) with less than 65% of reads mapped uniquely and an average depth of coverage less than 20 were flagged as failing alignment and removed from analysis. No samples were removed on the basis of failed sequence alignment.

**Variant Calling: PECaller[6]**

Variant calling was performed in a single batch using PECaller[6], which assumes multiple samples all done on the same technology will be available. Optimal PECaller[6] performance is achieved when at least 50 genomes are called in batch; 57 control genomes were included with the genomes from families 3206 and 3147 (63 genomes total). PECaller[6] was run with the default theta value of 0.001 and a 95% posterior probability for a genotype to be considered called. A posterior probability of less than 95% was considered a missing call. Calls were produced for the repeat-masked (unique) subset of the human Hg38 reference as reported by the University of California at Santa Cruz (UCSC) Genome Browser on July 1, 2015. The initial .snp file output from PECaller[6] was used in a subsequent step to merge SNP variant calls with INDEL variant calls, producing a final “merged” .snp file. This raw file was used for site and sample quality control.

**Whole-genome Sequence Quality Control**

Quality control was performed on a per-site and per-sample basis. The following metrics were used to flag and/or exclude samples and variant sites from QC and analysis, and were calculated using a custom QC pipeline consisting of multiple in-house-developed scripts, PLINK 1.9[5], R,[7] and Bystro[8]:

1. *Per-site QC: Missing call rate:* The missing call rates for variant sites were calculated as described above. Variants with a missing call rate greater than or equal to 10% were removed from subsequent QC *and* variant analysis.
2. *Sample Mixture Check:* Possible sample mixture was checked by calculate the ratios of minor allele homozygous calls to heterozygous calls. This number varies between call batches, and thus cannot be compared across different calling experiments. However, non-mixed samples within the same calling batch should exhibit similar ratios. The ratios were calculated using Bystro[8]. Any samples with ratios falling 3 SDs outside the mean were flagged for potential sample mixture, removed from analysis, and investigated. No samples were removed on the basis of possible sample mixture.
3. *Per-sample QC: Transition:Transversion Ratio:* Transition:transversion (Ti:Tv) ratios were calculated for each genome in the variant calling batch using a script developed in-house Bystro[8]. Based on population expectations, the Ti:Tv ratio for an individual genome is expected to be approximately 2.00, with a ratio of 2.04 representing a quality genome. The batch mean Ti:Tv ratio were calculated using Bystro[8]. The control genomes used in batch calling were previously validated for calling performance, therefore a *mean* Ti:Tv ratio less than 2.00 suggests a failed variant-calling experiment. As such, the entire sample batch is resubmitted for variant calling. Otherwise, any samples with a Ti:Tv ratio less than 2.00 were flagged and removed from subsequent QC. No samples were removed from analysis on the basis of Ti:Tv ratio.

4. *Per-sample QC: Silent:Replacement Ratio:* Silent:replacement (sil:rep) ratios were calculated for each genome in the variant calling batch using Bystro[8]. The expected sil:rep ratio for a single genome is expected for fall between 1.05 and 1.15, with 1.15 indicating a quality genome. The batch mean sil:rep ratio and standard deviation were calculated using Bystro[8]. A mean sil:rep

less than 1.05 suggested a failed variant-calling experiment and the sample batch was resubmitted for variant calling. Any samples with a sil:rep ratio less than 1.05 were flagged and removed from subsequent QC. No samples were removed from analysis on the basis of sil:rep ratio.

5. *Per-sample QC: Missing call rate:* The missing call rates for samples were calculated using PLINK[5]. The merged .snp file generated after the indel merging process was converted to a VCF [v4.0] format (snp_to_vcf2), the appropriate VCF headers were appended to file, and multiallelic variants were split using BCFtools 1.3[9], before the final BCF was loaded into PLINK[5]. The following flags were used during loading: *--bcf*, and *--keep-allele-order*. Per sample missing call rates were calculated using the *–missing* flag in PLINK[5], and the batch mean missing call rate and standard deviation was calculated using R[7]. A mean missing call rate greater than or equal to 3% indicated a failed variant-calling experiment and the sample batch was resubmitted for variant calling. Any sample(s) with a missing call rate greater than or equal to 3% were flagged and removed from subsequent QC. No samples in the current analysis were removed on the basis of low call rate.

1. *Sex Check:* PLINK was used to calculate the F coefficient estimates for the X chromosome and impute sex assignment for each sample in the batch. Before sex was inferred, the *--split-x* flag with the *hg38* modifier was used to identify pseudoautosomal regions of the X chromosomes for subsequent removal during the sex check. The default parameters for the *--check-sex* flag were used to infer sex. A sample with an F coefficient less than or equal to 0.2 was assigned as male, and a sample with an F coefficient greater than or equal to 0.9 was assigned as female. Any samples with an opposite sex assignment than indicated by the given pedigree were flagged and investigated for possible sample swapping or sample mixture. All samples’ inferred sex matched our expectation based on provided information.

2. *Relationship Inference:* Expected relationships between related samples of the batch were verified with PLINK. Variants were LD-pruned using the *--indep-pairwise* flag using 50, 5, and 0.2 for the variant count window size, variant count step size and $r^2$, respectively. The *--genome* flag was used to infer relationships (coefficient of relatedness; *r*) on this set of pruned SNPs. Among the control genomes there was a known parent-offspring relationship, which was used as a positive control, while the remaining control genomes were known to be unrelated. Any samples with an *r* estimate that indicated a degree of relationship (1$^{st}$, 2$^{nd}$, 3$^{rd}$, etc.) different from the expected degree of relationship were flagged and investigated for possible sample swapping, sample mixture, or misinformation. All samples' inferred relationships matched our expectations based on information provided by the families.

**Parental Origin Analysis**

Parental origin of the 3q29 deletion was determined for 12 trios --10 full trios and 2 trios for which only the child and mother's info was available -- using SNP array data. Briefly, using PLINK 1.9[5], SNPs located within the 3q29 deletion critical region (chr3: 196029182-197617792; hg38) were isolated for analysis. Mendelian errors (MEs) were called for these SNPs using PLINK's *--mendel* function with the *-duos* modifier to also call MEs for the mother-daughter pairs. The parent with the most mendelian errors was considered the parent of origin for the 3q29 deletion. Parental origin was determined using WGS data for two trios (3147 and 3206). Briefly, variants in the 3q29 critical region were called using PECaller. The variants with a sample minor allele frequency (MAF) less that 10% were filtered from this set of SNPs, and MEs were called. As in the SNP array analysis, the parent with the most MEs was considered the parent of origin for the 3q29 deletion

**Paternal Age Analysis**

Age of fathers at birth data for ~3 million U.S. births in 2018 (latest data available) were obtained from the National Center for Health Statistics (NCHS) (https://www.cdc.gov/nchs/index.htm). The means age of fathers in our 3q29 cohort was collected from self-reported data in conjunction with the Emory University 3q29 project (http://genome.emory.edu/3q29/) and compared to the U.S. average via a two-tailed two-sample t-test using R[7].

**Calculation of Recombination Rates**

Chromosome male and female recombination rates (cM/Mb) were obtained from the deCODE sex-specific maps[10]. The data from deCODE is presented as binned rates across separate chromosomes. As such, each binned recombination rate was weighted by the total basepairs of CNV contained within the respective bin (breakpoints cited in Coe et al, 2014[1]). Weighted binned rates were then averaged across the CNV interval.

**Linear Regression**

Parental origin data from multiple studies for each CNV locus mentioned above were combined into one sample size per locus. The $\log_e$-transformed combined male to female parental origin count ratios for each locus was regressed on the calculated $\log_e$-transformed average male to female recombination rate ratio for that locus' CNV interval using R[7]. Each locus was weighted based on its combined sample size. 95% confidence intervals for each locus were calculated using he following formula: $e^{\ln(M/F) \pm 1.96 \, \mathrm{sqrt}(1/N * (M/F + F/M))}$, where M = the total combined counts of male parental origins for each locus, F = the total combined count of female parental origins for each locus, and N = the combined sample size for each locus. Under the assumption that an NAHR event produces reciprocal deletion and duplication products, formation of both types of CNVs would be subject to the same biological forces. Thus, for each locus, duplications and deletions were treated equally and grouped under one locus.

We note that 17q11.2q12, a known genomic disorder locus associated with Charcot-Marie-Tooth disease type 1A (CMT1A; duplication) and hereditary neuropathy with liability to pressure palsies (HNPP; deletion), is mediated by NAHR, and thus applicable for inclusion in our analysis. However, subsequent research on the locus produced reports of a sex-dependent bias in both the mechanism for formation of the associated CNVs, and the resulting phenotype[11,12]. CNVs of paternal origin are generated via NAHR between homologous chromosomes during meiosis and are largely duplications (resulting in CMT1A), whereas CNVs of maternal origin are produced via intrachromosomal rearrangement between sister chromatids and result in equal numbers of deletions and duplications (resulting in CMT1A and HNPP). This is likely to cause a complex ascertainment bias and introduce a confounder associated with this locus. For this reason, we excluded the 17q11.2q12 locus from this study.

**Sensitivity Analysis**

A sensitivity analysis was conducted for the linear regression by iteratively running the linear model in R[7]. On each iteration one data point was removed from the model in order to identify potential influencing points. Results from the analysis are listed in Extended Data Table 3.

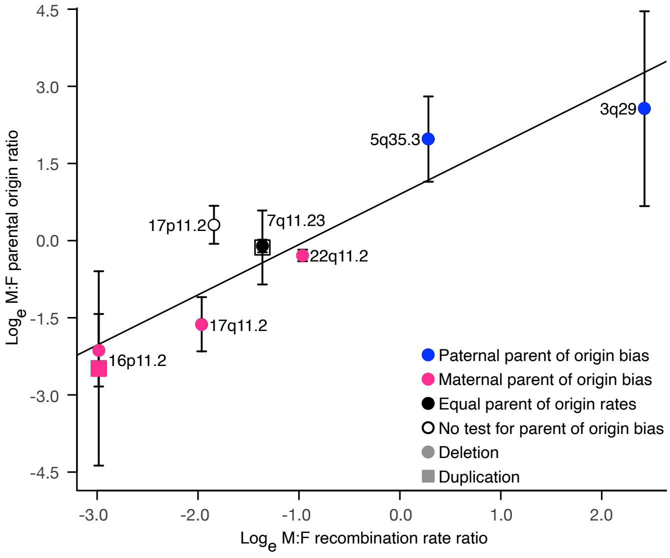


**Supplementary Figure Figure 1 | Recombination rates associate with parental origin bias.** $Log_e$-transformed male to female parental origin ratio regressed on $log_e$-transformed male to female recombination rate ratio. Recombination rates are associated with male to female parent origin ratios (multiple $R^2 = 0.8338$, $p=0.0005$, $\beta = 0.9780$, 95 % CI: (0.5877445,1.368187)). Curated parent of origin data from multiple published studies is separated into deletions and duplications at each locus. Data points are color coded by reported bias. Blue: loci with reported paternal biases, pink: loci with maternal biases, black: loci with equal male and female parental origin rates, white: loci where no test for parental origin bias conduced in literature. Circles and squares represent deletions and duplications, respectively. Error bars correspond to upper and lower bounds of 95% confidence interval. Note deletions (black circle) and duplications (white square) at 7q11.23 have similar male to female parental origin ratios and overlap on the plot.

**Supplementary Figure Table 1 l Demographic data for 3q29 cohort**

| Family ID | Subject ID | *Father ID | *Mother ID | Sex | Ethnicity | Race | Deletion Parental Origin | †Parent of Origin Age |
|---|---|---|---|---|---|---|---|---|
| 3156 | 834-3156-1031 | N/A | 834-3156-2096 | F | Non-Hispanic/Latino | White | Paternal | 24y |
| 3156 | 834-3156-2096 | | | F | Non-Hispanic/Latino | White | | |
| 3162 | 834-3162-1031 | N/A | 834-3162-2096 | F | Non-Hispanic/Latino | White | Paternal | 40y |
| 3162 | 834-3162-2096 | | | F | Non-Hispanic/Latino | White | | |
| 3164 | 834-3164-1001 | 834-3164-2046 | 834-3164-2096 | M | Non-Hispanic/Latino | White | Paternal | 32y |
| 3164 | 834-3164-2046 | | | M | Non-Hispanic/Latino | White | | |
| 3164 | 834-3164-2096 | | | F | Non-Hispanic/Latino | White | | |
| 3168 | 834-3168-1001 | 834-3168-2046 | 834-3168-2096 | M | Non-Hispanic/Latino | White | Paternal | 43y |
| 3168 | 834-3168-2046 | | | M | Non-Hispanic/Latino | White | | |
| 3168 | 834-3168-2096 | | | F | Non-Hispanic/Latino | White | | |
| 3181 | 834-3181-1001 | 834-3181-2046 | 834-3181-2096 | M | Hispanic/Latino | White | Paternal | 34y |
| 3181 | 834-3181-2046 | | | M | Hispanic/Latino | White | | |
| 3181 | 834-3181-2096 | | | F | Non-Hispanic/Latino | White | | |
| 3189 | 834-3189-1001 | 834-3189-2046 | 834-3189-2096 | M | Non-Hispanic/Latino | White | Paternal | 29y |
| 3189 | 834-3189-2046 | | | M | Non-Hispanic/Latino | White | | |
| 3189 | 834-3189-2096 | | | F | Non-Hispanic/Latino | White | | |
| 3226 | 834-3226-1031 | 834-3226-2046 | 834-3226-2096 | F | Non-Hispanic/Latino | White | Maternal | 41y |
| 3226 | 834-3226-2046 | | | M | Non-Hispanic/Latino | White | | |
| 3226 | 834-3226-2096 | | | F | Non-Hispanic/Latino | White | | |
| 3246 | 834-3246-1001 | 834-3246-2046 | 834-3246-2096 | M | Non-Hispanic/Latino | White | Paternal | 38y |
| 3246 | 834-3246-2046 | | | M | Non-Hispanic/Latino | White | | |
| 3246 | 834-3246-2096 | | | F | Non-Hispanic/Latino | White | | |
| 3257 | 834-3257-1031 | 834-3257-2046 | 834-3257-2096 | F | Non-Hispanic/Latino | White | Paternal | 38y |
| 3257 | 834-3257-2046 | | | M | Non-Hispanic/Latino | White | | |
| 3257 | 834-3257-2096 | | | F | Non-Hispanic/Latino | White | | |
| 3277 | 834-3277-1001 | 834-3277-2046 | 834-3277-2096 | M | Non-Hispanic/Latino | White | Paternal | 28y |
| 3277 | 834-3277-2046 | | | M | Non-Hispanic/Latino | White | | |
| 3277 | 834-3277-2096 | | | F | Non-Hispanic/Latino | White | | |
| 3396 | 834-3396-1031 | 834-3396-2046 | 834-3396-2096 | F | Non-Hispanic/Latino | White | Paternal | 43y |
| 3396 | 834-3396-2046 | | | M | Non-Hispanic/Latino | White | | |
| 3396 | 834-3396-2096 | | | F | Non-Hispanic/Latino | White | | |
| 3420 | 834-3420-1001 | 834-3420-2046 | 834-3420-2096 | M | Non-Hispanic/Latino | White | Paternal | 36y |
| 3420 | 834-3420-2046 | | | M | Non-Hispanic/Latino | White | | |
| 3420 | 834-3420-2096 | | | F | Non-Hispanic/Latino | White | | |
| 3206 | 834-3206-1031 | 834-3206-2096 | 834-3206-2046 | F | Non-Hispanic/Latino | White | Paternal | 32y |
| 3206 | 834-3206-2046 | | | M | Non-Hispanic/Latino | White | | |
| 3206 | 834-3206-2096 | | | F | Non-Hispanic/Latino | White | | |
| 3147 | 834-3147-1001 | 834-3147-2046 | 834-3147-2096 | M | Non-Hispanic/Latino | White | Paternal | 29y |
| 3147 | 834-3147-2046 | | | M | Non-Hispanic/Latino | White | | |
| 3147 | 834-3147-2096 | | | F | Non-Hispanic/Latino | White | | |

Demographic data is self-reported.
*N/A indicates information is not available. Grandparental samples were not collected.
†Age of parent of origin is corresponds to parent's at birth of affected child.

**Supplementary Figure Table 2 | Extended data for linear regression analysis with deletions and duplications combined**

| Locus | *CNV Type(s) | †BED Coordinates[10] | Sample Size | ‡Paternal Origin Counts | ‡Maternal Origin Counts | M:F Origin Ratio | §Male Recombination Rate (cM/Mb)[54] | §Female Recombination Rate (cM/Mb)[54] | $Log_e$ M:F Recombination Rate Ratio |
|---|---|---|---|---|---|---|---|---|---|
| 3q29 | Del | chr3:195988732-197628732 | 14 | 13 | 1 | 13 | 3.14154034 | 0.2782546 | 2.4239321 |
| 5q35.3 | Del | chr5:176290391-177630393 | 41 | 36 | 5 | 7.2 | 1.29955355 | 0.9794135 | 0.2828221 |
| 7q11.23 | Del/Dup | chr7:73328061-74727726 | 530 | 251 | 279 | 0.899641577 | 0.4940131 | 1.9286881 | -1.3620333 |
| 16p11.2 | Del/Dup | chr16:29641178-30191178 | 79 | 8 | 71 | 0.112676056 | 0.06553698 | 1.2949196 | -2.9835893 |
| 17p11.2 | Del | chr17:16805961-20576095 | 59 | 34 | 25 | 1.36 | 0.1888066 | 1.1911597 | -1.8419594 |
| 17q11.2 | Del | chr17:30838856-31888868 | 73 | 12 | 61 | 0.196721311 | 0.26024285 | 1.8544277 | -1.9637162 |
| 22q11.2 | Del | chr22:19032487-20302477 | 642 | 275 | 367 | 0.749318801 | 1.37677793 | 3.6065407 | -0.9630031 |

*Data from published studies (See Data S1) include parental origin reports for deletions and duplications at 7q11.23 and 16p11.2.
†Hg38 BED coordinates as cited in Coe et al., 2014[10]
‡Combined raw numbers of paternal and maternal origins from published studies
§Average recombination rates. Calculated from sex-specific rates published in Halldorsson et al., 2019[54]

**Supplementary Figure Table 3 l Sensitivity analysis results for linear regression analysis with deletions and duplications combined**

| Locus Removed | CNV Type | *$R^2$ | p-value | β |
|---|---|---|---|---|
| None | Deletion/Duplication | 0.8344 | 0.004 | 0.9728 |
| 3q29 | Deletion | 0.8141 | 0.014 | 1.1123 |
| 5q35.3 | Deletion | 0.8245 | 0.012 | 0.8883 |
| 7q11.23 | Deletion/Duplication | 0.8719 | 0.006 | 0.9856 |
| 16p11.2 | Deletion/Duplication | 0.7227 | 0.032 | 0.8979 |
| 17p11.2 | Deletion | 0.8791 | 0.006 | 0.9977 |
| 17q11.2 | Deletion | 0.8441 | 0.010 | 0.9362 |
| 22q11.2 | Deletion | 0.8693 | 0.007 | 1.0221 |

*Multiple $R^2$ value as reported by R.
Our sensitivity analysis indicates the estimates of the variation in parental bias explained by recombination is not influenced by any particular data point.

**Supplementary Figure Table 4 I Extended data for linear regression analysis with deletions and duplications separated**

| Locus | CNV Type | *BED Coordinates[10] | Sample Size | †Paternal Origin Counts | †Maternal Origin Counts | M:F Origin Ratio | ‡Male Recombination Rate (cM/Mb)[54] | ‡Female Recombination Rate (cM/Mb)[54] | $Log_e$ M:F Recombination Rate Ratio |
|---|---|---|---|---|---|---|---|---|---|
| 3q29 | Del | chr3:195988732-197628732 | 14 | 13 | 1 | 13 | 3.14154034 | 0.2782546 | 2.4239321 |
| 5q35.3 | Del | chr5:176290391-177630393 | 41 | 36 | 5 | 7.2 | 1.29955355 | 0.9794135 | 0.2828221 |
| 7q11.23 | Del | chr7:73328061-74727726 | 515 | 244 | 271 | 0.900369 | 0.4940131 | 1.9286881 | -1.3620333 |
| 7q11.23 | Dup | chr7:73328061-74727726 | 15 | 7 | 8 | 0.875 | 0.4940131 | 1.9286881 | -1.3620333 |
| 16p11.2 | Del | chr16:29641178-30191178 | 66 | 7 | 59 | 0.1186441 | 0.06553698 | 1.2949196 | -2.9835893 |
| 16p11.2 | Dup | chr16:29641178-30191178 | 13 | 1 | 12 | 0.08333333 | 0.06553698 | 1.2949196 | -2.9835893 |
| 17p11.2 | Del | chr17:16805961-20576095 | 59 | 34 | 25 | 1.36 | 0.1888066 | 1.1911597 | -1.8419594 |
| 17q11.2 | Del | chr17:30838856-31888868 | 73 | 12 | 61 | 0.1967213 | 0.26024285 | 1.8544277 | -1.9637162 |
| 22q11.2 | Del | chr22:19032487-20302477 | 642 | 275 | 367 | 0.7493188 | 1.37677793 | 3.6065407 | -0.9630031 |

*Hg38 BED coordinates as cited in Coe et al., 2014[10].
†Combined raw numbers of paternal and maternal origins from published studies.
‡ Average recombination rates. Calculated from sex-specific rates published in Halldorsson et al., 2019[54].
